# Chitinase 3-like-1 Stimulates PD-L1 and Other Immune Checkpoint Inhibitors

**DOI:** 10.1101/2021.01.15.426788

**Authors:** Bing Ma, Bedia Akosman, Suchitra Kamle, Chang-Min Lee, Ja Seok Koo, Chun Geun Lee, Jack A. Elias

## Abstract

PD-1 and its ligand PD-L1 are major mediators of tumor-induced immunosuppression. Chitinase 3-like-1 (Chi3l1) is induced in many cancers where it portends a poor prognosis and contributes to tumor metastasis. Here we demonstrate that Chi3l1 regulates the expression of PD-L1, PD-L2, PD-1 and LAG3 in melanoma lung metastasis. Chi3l1 stimulates macrophage PD-L1 expression and mediates optimal IFN-γ-stimulated PD-L1 expression via IL-13Rα2. We also demonstrate that RIG-like helicase innate immune activation suppresses Chi3l1, PD-L1, LAG3 and pulmonary metastasis. At least additive antitumor responses were seen in metastasis models treated simultaneously with individual antibodies against PD-1 and Chi3l1. At least additive cytotoxic T cell-induced tumor cell death was also seen in co-cultures of T and tumor cells treated with antibodies that target Chi3l1 and PD-1. Thus, Chi3l1 contributes to pulmonary metastasis by stimulating the PD1-PD-L1 axis and other checkpoint molecules. The simultaneous targeting of Chi3l1 and the PD-1-PD-L1 axis, represents a promising therapeutic strategy for pulmonary metastasis.

## INTRODUCTION

One of the most exciting recent discoveries in cancer is the appreciation that evasion of the immune system and suppression of neoantigen-induced T cell responses are essential for cancer development, progression and resistance to treatment (1–3). Studies of these responses have highlighted essential immune checkpoints (ICP) that are regulated by inhibitory receptors and their ligands such as programed death (PD)-1 and its ligands PD-L1 and PD-L2; cytotoxic T lymphocyte antigen 4 (CTLA4) and its ligands B7.1/B7.2; and lymphocyte activation gene 3 protein (LAG3) and its ligand HLA class II (4). These advances have led to the development of immune checkpoint molecule blocking antibodies against PD-1, PD-L1, and CTLA4 which have proven to be impressively successful therapeutics for a number of malignancies (2, 4). This is particularly important in the lung because recent studies demonstrated that ICP antibodies can be useful therapeutics in lung cancer (5–7). In these and other studies of ICP inhibitor-based therapeutics, some patients respond impressively while others do not. In many cases responsiveness to interventions in the PD-1-PD-L1 axis correlate with the expression of molecules like PD-L1 (5–7). However, the moieties that regulate the expression of immune checkpoint molecules and the complex mechanisms that they employ (8), have not been adequately identified.

Chitinase 3-like-1 (Chi3l1; also called YKL-40), the prototypic chitinase-like protein (CLP), was originally discovered in mouse breast cancer cells (9). It is now known to be expressed by a variety of cells including macrophages, neutrophils and epithelial cells and is stimulated by a number of mediators including IL-13, IL-6, IL-1β, and IFN-γ (10–13). Studies from our laboratory and others have also demonstrated that Chi3l1 is a multifaceted moiety that inhibits cell death (apoptosis and pyroptosis), stimulates Th2 inflammation and M2 macrophage differentiation, inhibits oxidant injury, controls inflammasome and caspase activation, regulates TGF-β1 elaboration, contributes to antibacterial responses and activates MAP Kinase (MAPK), Akt/Protein Kinase B and Wnt/β-catenin signaling (10, 14–17). They also demonstrated that many of these responses are mediated by a multimeric receptor called the chitosome that contains an IL-13Rα2 alpha subunit and a TMEM219 (TMEM) β subunit (15, 18). In keeping with these diverse sources and stimuli, elevated levels of Chi3l1 have been noted in a wide variety of diseases characterized by inflammation, fibrosis and tissue remodeling (11, 19–22). However, the roles of Chi3l1 in these diverse diseases have not been fully defined.

Recent studies demonstrated that the levels of circulating Chi3l1 are increased in many malignancies including cancers of the prostate, colon, rectum, ovary, kidney, breast, glioblastomas and malignant melanoma (23–35). In these diseases, the levels of Chi3l1 frequently correlate directly with disease progression and inversely with disease-free interval and survival (23–35). This is particularly striking in lung cancer where the serum and tissue levels of Chi3l1 are impressively increased and correlate with adverse outcomes (36–39). Chi3l1 may also play a particularly important role in pulmonary metastasis where studies from our lab and others have demonstrated that Chi3l1 induction is required for the generation of a metastasis permissive pulmonary microenvironment (40) and metastatic spread can be inhibited via RIG-like helicase (RLH) innate immune inhibition of Chi3l1 elaboration (41). However, the mechanism(s) by which Chi3l1 contributes to tumor initiation and spread have not been adequately defined and the degree to which Chi3l1 mediates its tumorigenic effects via activating ICP pathways such as the PD1-PDL1 axis has not been addressed. In addition, although recent studies reported that RLH activation is critical for responsiveness to immune checkpoint blockade (42), the possibility that this responsiveness is mediated by RLH immune inhibition of Chi3l1 has not been considered.

We hypothesized that Chi3l1 contributes to pulmonary metastasis via the regulation of immune checkpoint molecules. To address this hypothesis, studies were undertaken to determine if Chi3l1 regulates the expression of components of the PD1-PDL1 axis and or LAG3. *In vivo* studies demonstrated that PD-L1, PD-L2, PD-1 and LAG 3 are induced by melanoma metastasis via Chi3l1-dependent mechanisms and that transgenic Chi3l1 stimulates these checkpoint inhibitors. *In vitro* studies demonstrated that, Chi3l1 stimulates macrophage PD-L1 via a mechanism(s) that utilizes IL-13 Receptor α2 (IL-13Rα2) and that optimal gamma interferon (IFN-γ) stimulation of macrophage PD-L1 requires Chi3l1. Our studies also demonstrated that RLH innate immune activation suppressed Chi3l1, PD-L1 and pulmonary metastasis. From a therapeutic perspective, at least additive antitumor effects were seen in metastasis models treated simultaneously with individual antibodies against PD1 and Chi3l1. Synergistic cytotoxic T cell (CTL)-induced tumor cell death was seen in T cell-tumor co-cultures treated with bispecific antibodies that simultaneously target Chi3l1 and PD1. These studies demonstrate that Chi3l1 contributes to the development and or progression of metastatic tumors in the lung via the regulation of PD1-PDL1 axis and other immune checkpoint molecules. They also strongly support the concept that the simultaneous inhibition of Chi3l1 and components of the PD-1-PD-L1 axis with monospecific antibodies represents is a novel and attractive therapeutic strategy for lung metastasis.

## RESULTS

### Pulmonary melanoma metastasis stimulates PD-L1

Studies were undertaken to determine if the expression of immune checkpoint molecules (ICPs) was altered by the metastatic spread of melanoma to the lung. In these experiments, 8 weeks old C57BL/6 mice were challenged with freshly prepared B16-F10 (B16) melanoma cells or vehicle control and the pulmonary expression of ICPs was evaluated 2 weeks later. These studies demonstrated that melanoma metastasis was associated with significantly increased expression of multiple ICP moieties including PD-1, PD-L1, and PD-L2. (Fig. 1A and Fig. S1). This induction was not specific for the PD-1/PD-L1/PD-L2 axis because LAG3 was also induced (Fig. S1). Among these events, the induction of PD-L1 mRNA and protein were particularly prominent in comparisons of lungs from B16-F10 (B16) and vehicle challenged mice (Fig. 1, A and B). FACS analysis demonstrated that this enhanced expression of PD-L1 was seen in a variety of cells including airway and alveolar epithelial cells (CC10 and SPC positive cells respectively), CD3 positive T cells and CD68 positive macrophages (Fig. 1C). Double label immunohistochemical staining using antibodies against cell specific markers and PD-L1 further reinforced the enhanced expression of alveolar epithelial cell and macrophage PD-L1 (Fig. 1, D and E). These studies demonstrate that pulmonary melanoma metastasis is associated with significantly enhanced expression and accumulation of ICPIs including PD-1/PD-L1 and PD-L2.

**Fig. 1.**
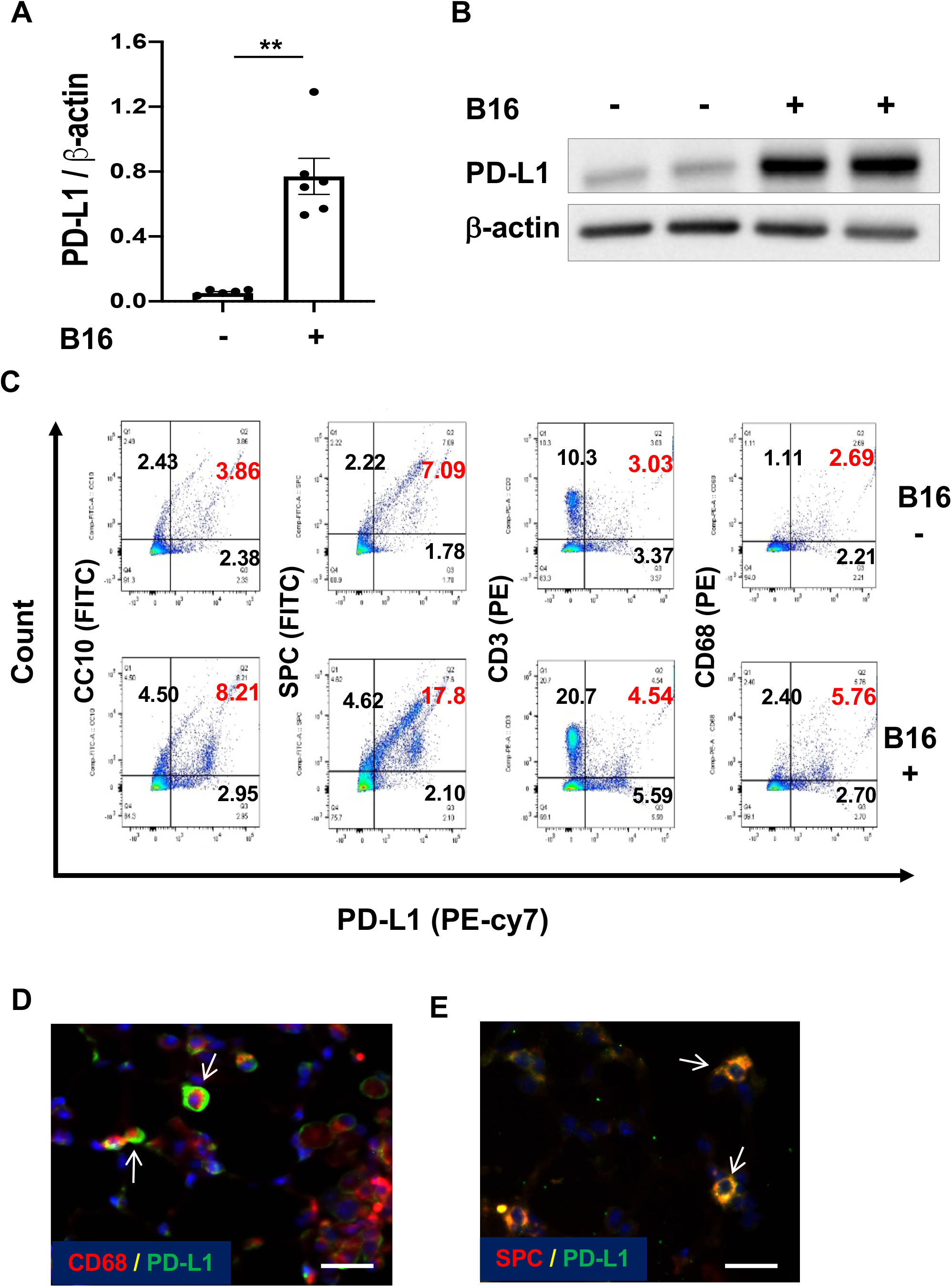
Pulmonary melanoma metastasis stimulates PD-L1. 8 week-old WT mice were given B16-F10 (B16) melanoma cells or control vehicle (PBS) and evaluated 2 weeks later. (A) Realtime RT-PCR (RT-PCR) was used to quantitate the levels of mRNA encoding PD-L1 in the lungs from mice treated intravenously with PBS (B16 -) or B16 cells (B16 +). Each dot represents an evaluation in an individual animal. (B) Western blot evaluations of PD-L1 accumulation in lungs from mice treated with PBS (B16-) or B16 cells (B16 +). (C) FACS evaluations quantitating the accumulation of PD-L1 in cell populations from lungs of mice treated with B16 cells (B16+) or vehicle control (B16-). These evaluations used specific markers of airway epithelial cells (CC10), alveolar epithelial cells (surfactant apoprotein C; SPC), T cells (CD3), and macrophages (CD68). (D and E) Representative double label fluorescent immunohistochemistry (IHC) evaluations comparing lungs from mice challenged with vehicle or B16 melanoma cells using cell-specific markers (alveolar epithelial cells; surfactant apoprotein C; SPC), macrophages (CD68) (red) and anti-(α)-PD-L1 (green). The arrows highlight cells that stain with both antibodies. The values in panel A represent the mean± SEM of the noted evaluations represented by the individual dots. Panels B, C, D and E are representative of a minimum of 2 similar evaluations. **P<0.01. Scale bars = 50 μm

### Chi3l1 plays a critical role in B16 melanoma metastasis stimulation of pulmonary PD-L1

Studies were next undertaken to define the potential role(s) of Chi3l1 in the induction of PD-L1 during the course of B16 cell pulmonary metastasis. As noted above, the levels of mRNA encoding PD-L1 were significantly increased in lungs from melanoma challenged mice compared to lungs from PBS controls (Fig.2A). This induction was significantly reduced in the lungs from mice with null mutations of Chi3l1 (Fig. 2A). Accordingly, PD-L1 protein accumulation was also increased in the lungs from mice challenged with melanocytes compared to vehicle controls and this induction was significantly decreased in lungs from Chi3l1 null animals (Fig. 2B). B16 cell stimulation of PD-L1 expression was also significantly diminished in lungs from mice treated with monoclonal anti-Chi3l1 antibody (called FRG antibody) (Fig. 2C). In accord with these findings, the ability of B16 cells to induce the accumulation of PD-L1 protein was also diminished by treatment with anti-Chi3l1 antibody (Fig. 2D). These studies were reinforced by double label immunohistochemical evaluations which highlighted the impressive induction of PD-L1 in CD68 + macrophages in lungs from B16 challenged mice and the decrease in PD-L1 accumulation in similarly challenged Chi3l1 null animals (Fig. 2E). The importance of Chi3l1 in these inductive events was not unique to PD-L1 because null mutations of Chi3l1 and treatment with anti-Chi3l1 had similar effects on PD-1 and LAG3, respectively (Fig. S2). When viewed in combination, these studies demonstrated that Chi3l1 plays an essential role in melanoma stimulation of PD-L1 and other ICPs.

**Fig. 2.**
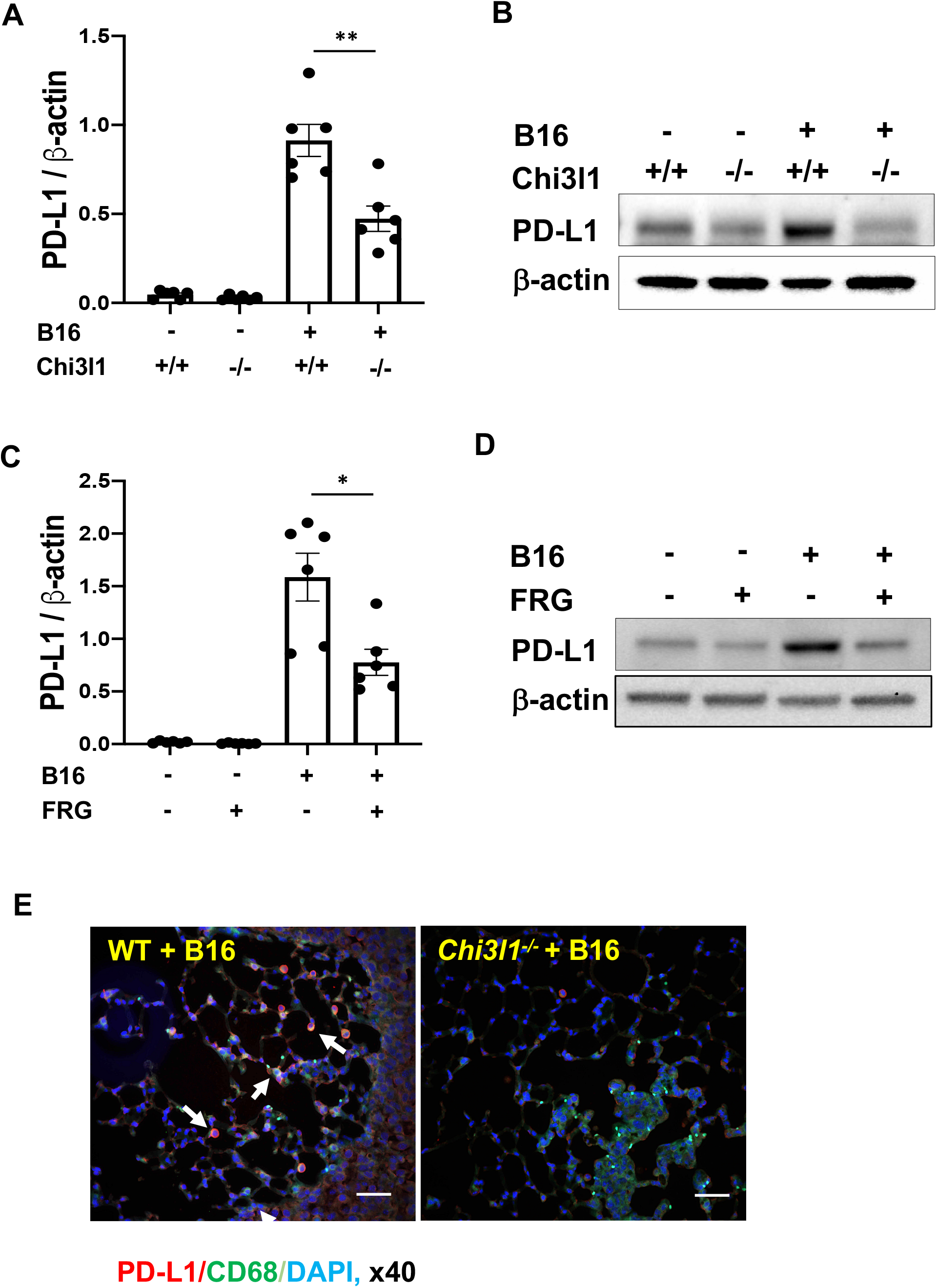
Chi3l1 plays a critical role in B16 melanoma stimulation of pulmonary PD-L1. 8 week-old WT (+/+) and Chi3l1 null (-/-) (Chi3l1^-/-^) mice were given B16 melanoma cells or vehicle control. They were then treated with an anti-Chi3l1 antibody (FRG) or isotype control antibodies and PD-L1 expression was evaluated 2 weeks later. (A) RT-PCR was used to quantitate the levels of mRNA encoding PD-L1 in the lungs from mice treated intravenously with PBS vehicle (B16 -) or B16 cells (B16 +). Wild type (WT) (Chi3l1 +/+) and Chi3l1 null (Chi3l1 -/-) mice were employed. Each dot represents an evaluation in an individual animal. (B) Western blot evaluations of PD-L1 accumulation in lungs from WT (Chi3l1^+/+^) and Chi3l1 null (Chi3l1^-/-^) mice treated with vehicle (B16-) or B16 cells (B16 +). (C) RT-PCR was used to quantitate the levels of mRNA encoding PD-L1 in the lungs from mice treated intravenously with vehicle (B16 -) or B16 cells (B16 +). The mice were treated with antibodies against Ch3l1 (FRG+) or isotype control antibodies (FRG -). Each dot represents an evaluation in an individual animal. (D) Western blot evaluations of PD-L1 accumulation in lungs from WT mice that were given control vehicle (B16-) or B16 cells (B16 +) and treated with antibodies against anti-Chi3l1 (FRG+) or isotype control (FRG -) antibodies. (E) Double label IHC comparison of lungs from WT and Chi3l1^-/-^ mice challenged with B16 melanoma cells using a macrophage-specific marker (CD68; green) and anti-PD-L1 antibodies (red). The plotted values in panels A & C represent the mean ± SEM of the noted evaluations represented by the individual dots. Panels B, D and E are representative of a minimum of 2 similar evaluations. *P<0.05, **P<0.01. Scale bars = 100μm.

### Transgenic Chi3l1 stimulates PD-L1 in the normal lung

The studies noted above demonstrate that pulmonary melanoma metastasis stimulate PD-L1 and other ICPs and that Chi3l1 plays a critical role in these inductive events. However, because null mutations of Chi3l1 and treatment with anti-Chi3l1 decrease metastatic spread (40, 41), the studies do not determine if the decreased induction of PD-L1 and other ICPs that is seen in Chi3l1 null mutant mice or mice treated with anti-Chi3l1 are due to the direct effects of Chi3l1 or the importance of Chi3l1 in melanoma metastatic spread. To address this issue, studies were undertaken to determine if Chi3l1 stimulates pulmonary PD-L1 in the absence of B16 cell administration. In these experiments, we compared the expression and accumulation of PD-L1 in lungs from WT mice and transgenic mice (Tg) in which Chi3l1 is overexpressed in a lung-specific manner. These experiments demonstrated that Chi3l1 is a potent stimulator of PD-L1 mRNA and protein in lungs from Tg mice when compared to WT controls (Fig. 3, A and B). FACS evaluations also demonstrated that transgenic Chi3l1 stimulated PD-L1 expression in a variety of cells including epithelial cells and macrophages (Fig. 3, C-F). This effect was not PD-L1-specific because transgenic Chi3l1 also stimulated PD-1, PD-L2 and LAG3 (Fig. S3). These studies demonstrate that the stimulation of PD-L1 and other ICPs in pulmonary melanoma metastasis is mediated, at least in part, by a tumor-independent effect of Chi3l1.

**Fig. 3.**
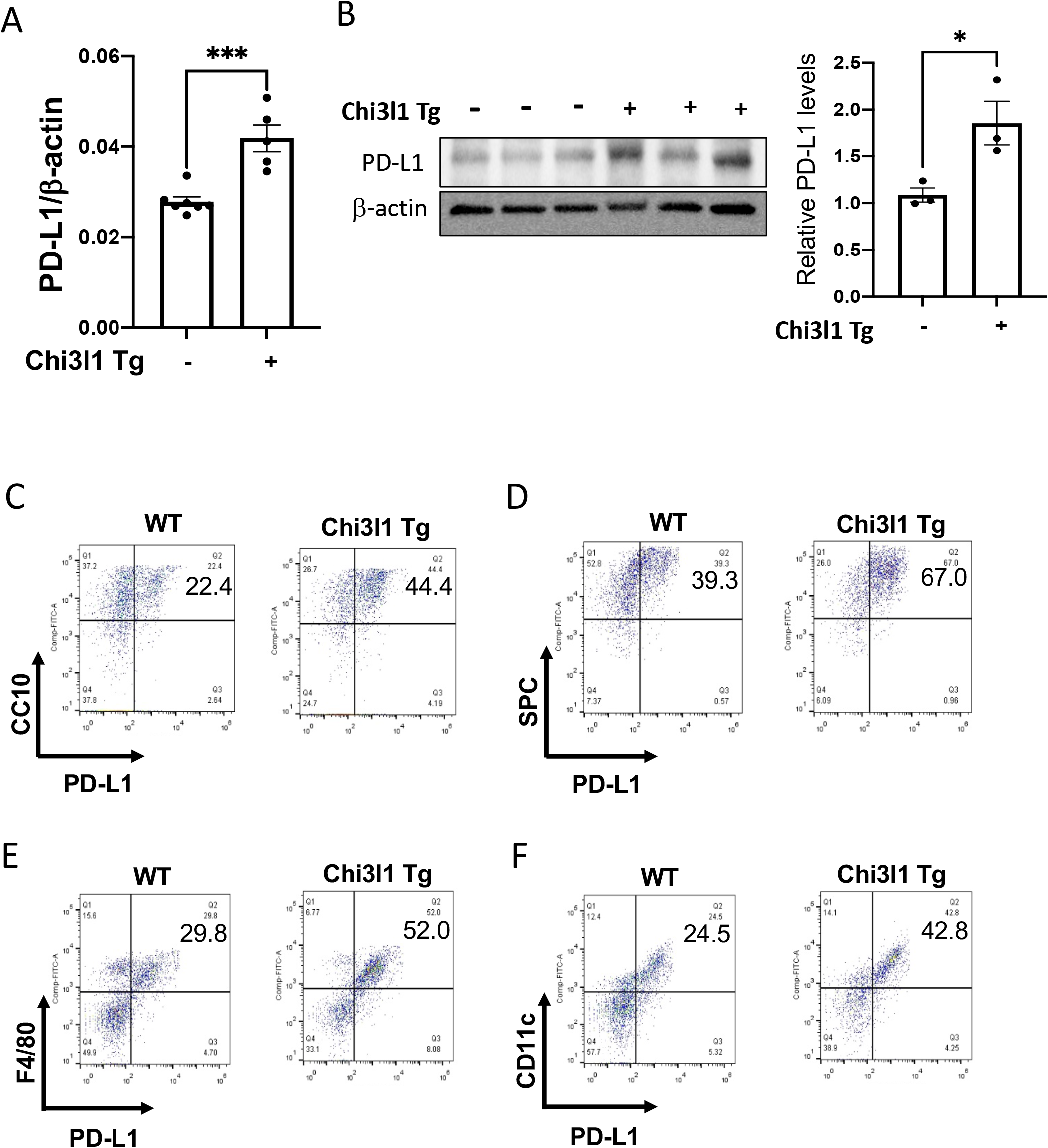
Transgenic Chi3l1 stimulates PD-L1 in the normal lung. 8 week-old WT (-) and Chi3l1 Tg (+) mice were used to evaluate the expression and accumulation of PD-L1 in the lung. (A) RT-PCR was used to quantitate the levels of mRNA encoding PD-L1 in the lungs from WT mice (Chi3l1 Tg-) and mice in which Chi3l1 was overexpressed in the lung in a transgenic manner (Chi3l1 Tg +). Each dot represents the evaluation in an individual animal. (B) Western blot evaluation of PD-L1 accumulation in lungs from WT (Chi3l1 Tg-) and Chi3l1 transgenic (Chi3l1 Tg+) mice and densitometric quantitation of relative expression of PD-L1. (C) FACS evaluation quantitating the accumulation of PD-L1 in cells populations from lungs from WT and Chi3l1 Tg + mice. These evaluations used cell-specific markers of airway epithelial cells (CC10), alveolar epithelial cells (surfactant apoprotein C; SPC), dendritic cells (CD11 c), and macrophages (F4/80). The values in panel A represent the mean ± SEM of the noted evaluations represented by the individual dots. Panels B-F are representative of a minimum of 2 similar evaluations. *P<0.05.

### Chi3l1 stimulates pulmonary macrophage PD-L1

Because macrophages were one of the major cells expressing PD-L1 in response to transgenic Chi3l1, studies were undertaken to further understand the mechanism(s) of this inductive event. As can be seen in Fig. 4A, the recovery of CD11b+ PD-L1 + cells was increased in lungs from Chi3l1 Tg (YKL-40 Tg) mice and decreased in lungs from Chi3l1 null animals when compared to WT controls. In addition, the stimulatory effect of transgenic Chi3l1 (YKL-40) on CD68+ macrophages was significantly decreased in mice treated with anti-Chi3l1 antibody (FRG) compared to animals treated with an isotype antibody control (Fig. 4B). We also compared the levels of mRNA encoding PD-L1 and PD-L1 protein in bone marrow-derived macrophages from WT mice treated with rChi3l1 or vehicle control. In these experiments rChi3l1 was a potent, dose dependent stimulator of PD-L1 mRNA and protein (Fig. 4, C and D). PD-L2 was similarly stimulated (Fig. 4E). These studies demonstrate that Chi3l1 stimulates the expression and accumulation of PD-L1 mRNA and protein in macrophages *in vivo* and *in vitro.*

**Fig. 4.**
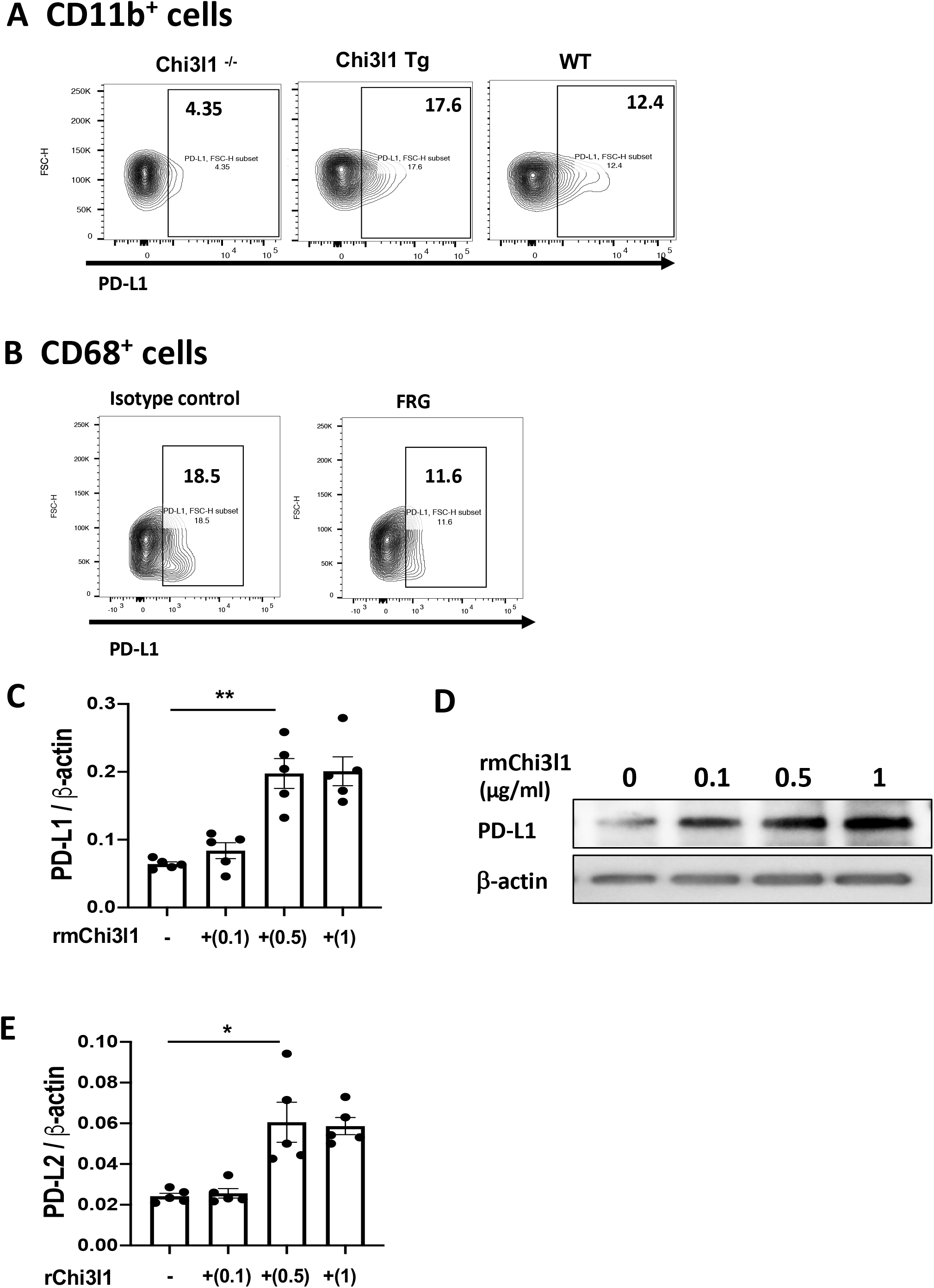
Chi3l1 stimulates pulmonary macrophage PD-L1. 8 week-old WT, Chi3l1 Tg (+) and Chi3l1 null (Chi3l1^-/-^) mice were used to evaluate macrophage lineage cell PD-L1 in the lung. (A) FACS evaluations comparing the PD-L1 on CD11b+ cells isolated from lungs from Chi3l1 null (Chi3l1^-/-^), Chi3l1 Tg, and WT mice. (B) FACS evaluations comparing the PD-L1 on CD68+ cells isolated from lungs from WT mice treated with anti-Chi3l1 antibody (FRG) or its isotype control. (C) RT-PCR was used to quantitate the levels of mRNA encoding PD-L1 in bone marrow-derived macrophages (BMDM) from WT mice after stimulation with the noted concentrations of recombinant murine (rm) Chi3l1 (μg/ml) or vehicle control. Each dot represents an evaluation performed using cells from an individual animal. (D) Western blot evaluations of PD-L1 accumulation in bone marrow-derived macrophages from WT mice treated with the noted concentrations of rmChi3l1 or vehicle control. (E) RT-PCR was used to quantitate the levels of mRNA encoding PD-L2 in bone marrow derived macrophages (BMDM) from WT mice after stimulation with the noted concentrations of rmChi3l1 (μg/ml) or vehicle control. Each dot represents an evaluation performed using cells from an individual animal. Panels A, B, and D are representative of a minimum of 2 similar evaluations. The values in panels C & E represent the mean ± SEM of the noted evaluations represented by the individual dots. *P<0.05, **P<0.01.

### Chi3l1 plays a critical role in IFN-γ-stimulation of macrophage PD-L1

Previous studies reported that IFN-γ is a potent stimulator of PD-L1 in macrophages and other immune cells (43, 44). Thus, studies were undertaken to define the role(s) of Chi3l1 in IFN-γ-stimulation of PD-L1. In these experiments, bone marrow (BM)-derived macrophages were obtained from WT and Chi3l1 null mutant mice and were incubated with recombinant (r) IFN-γ or vehicle control for up to 72 hrs. As shown in Fig 5A, rIFN-γ increased the levels of mRNA encoding PD-L1 in a time- and dose-dependent manner in WT cells (Fig. 5, A & B). rIFN-γ also stimulated macrophage Chi3l1 mRNA and protein expression in dose- and time-dependent manner (Fig. 5, C&D). Interestingly, the levels of rIFN-γ-stimulated PD-L1 protein were significantly decreased in Chi3l1 null macrophages when compared to macrophages from WT animals (Fig. 5E). A similar decrease in PD-L1 accumulation was seen in FACS evaluations of IFN-γ stimulated cells from Chi3l1 null mice versus WT controls (Fig. 5F). IFN-γ also stimulated macrophage PD-L2 in a Chi3l1-dependent manner (Fig. S4) Null mutations of IL-13Rα2 also significantly reduced IFN-γ-stimulation of macrophage PD-L1 (Fig. 5G). In contrast, significant changes were not noted in the cells with null mutations of or cells treated with inhibitors of other putative Chi3l1 receptors or interacting partners such as TMEM219, galectin-3 or CRTH2 (Fig. 5G) (18, 20, 45). In combination, these studies demonstrate that Chi3l1 and IL-13Rα2 play critical roles in IFN-γ stimulation of macrophage PD-L1.

**Fig. 5.**
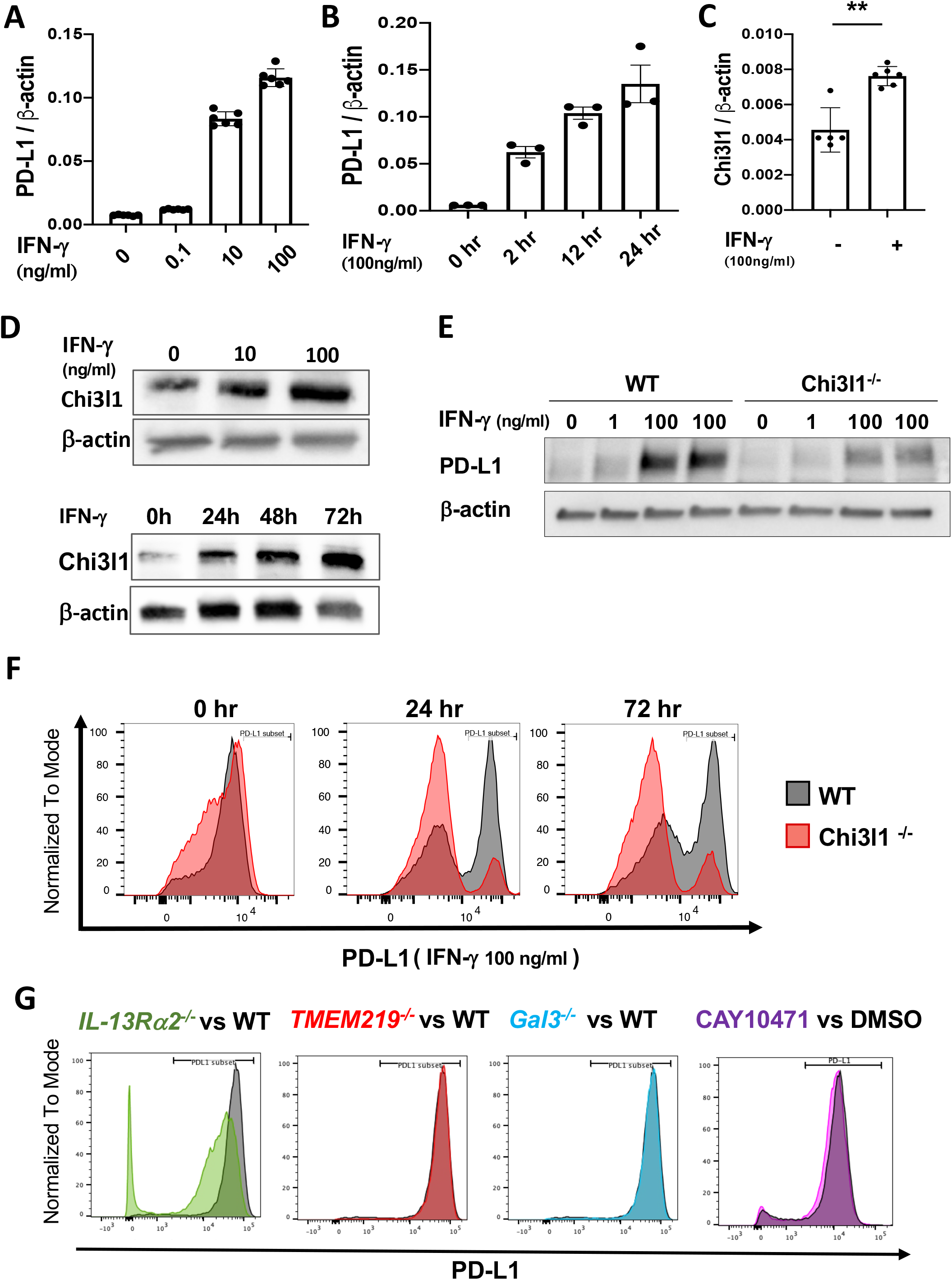
IFN-γ-stimulates macrophage PD-L1 via a Chi3l1-dependent mechanism. Bone marrow derived macrophages (BMDM) prepared from 6-8 week-old WT, Chi3l1^-/-^, IL-13Rα2^-/-^, TMEM219^-/-^, and Galectin 3 null (Gal-3^-/-^) mice and were used to evaluate the importance of Chi3l1 in rIFN-γ stimulation of PD-L1. (A-B) Dose and time dependency of IFN-γ-stimulation of PD-L1 mRNA in BMDM. BMDM from WT mice were incubated with the noted concentrations of rIFN-γ for the noted periods of time. (C) RT-PCR evaluation of the expression of Chi3l1 in BMDM after stimulation with recombinant IFN-γ for 24 hr. (D) Western blot evaluations of the dose and time-dependency of IFN-γ stimulation of macrophage Chi3l1 accumulation. (E) Western blot evaluations of rIFN-γ-stimulated PD-L1 accumulation in BMDM prepared from WT and Chi3l1^-/-^ mice. (F) FACS evaluations of the ability of IFN-γ-to stimulate PD-L1 in BMDM prepared from WT and Chi3l1^-/-^ mice. (G) FACS evaluations of the ability of IFN-γ-to stimulate PD-L1 accumulation in BMDM prepared from WT, IL-13Rα2^-/-^, TMEM219^-/-^ and Gal3^-/-^ mice and WT BMDM treated with vehicle (5% DMSO) or the selective CRTH2 inhibitor (Cay10471, 20μg/ml in 5%DMSO). The values in panels A-C represent the mean ± SEM of the evaluations represented by the individual dots. Panels D-G are representative of a minimum of 2 similar evaluations. **P<0.01.

### Rig-like helicase (RLH) activation inhibits Chi3l1 and PD-L1

We previously demonstrated that Poly(I:C), a strong activator of retinoic acid inducible gene I (RIG-I) and the RIG-like helicase (RLH) innate immune response, prominently inhibits Chi3l1 expression and melanoma lung metastasis (41). Recent studies also demonstrated that RIG-I activation and mitochondrial antiviral signaling molecule (MAVS) are essential for many immune checkpoint inhibitor blockade-induced anti-tumor responses (42). Thus, studies were undertaken to determine if RLH activation with Poly(I:C) altered the ability of Chi3l1 to stimulate PD-L1. As shown in Figures 6A and 6B, Poly(I:C) ameliorated melanoma-stimulated PD-L1 mRNA expression and inhibited B16 cell stimulation of Chi3l1 and PD-L1 protein accumulation (Fig. 6B). Poly(I:C) similarly inhibited LAG3 (Fig. 6C). FACS analysis also demonstrated that macrophage expression of PD-L1 was prominently reduced by poly(I:C) treatment (Fig. 6D). The suppressive effects of Poly(I:C) was at least partially dependent on Chi3l1 because the transgenic overexpression of Chi3l1 using a promoter that is not regulated by the RLH pathway ameliorated Poly(I:C)-induced inhibition of PD-L1 and LAG3, enhanced B16-stimulated expression of PD-L1 and LAG3 (Fig. 6, E and F). These studies highlight the ability of RLH activation to inhibit Chi3l1 and, in turn, inhibit PD-L1 and LAG3.

**Fig. 6.**
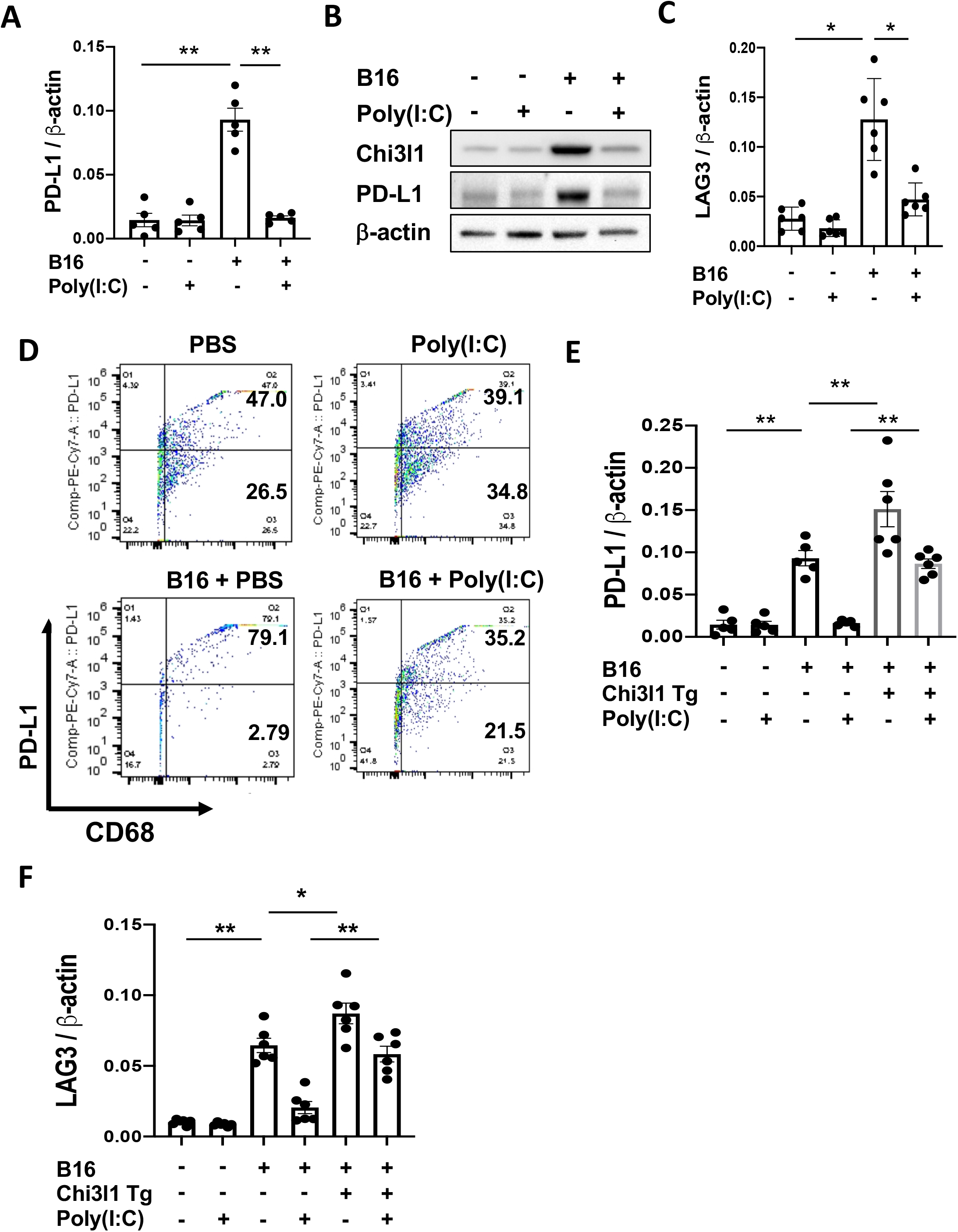
Rig like helicase (RLH) activation inhibits the induction of Chi3l1 and PD-L1. WT mice were given B16-F10 (B16) melanoma cells or PBS control and treated with Poly(I:C) or its vehicle control and evaluated 2 weeks later. (A and B) RT-PCR and Western evaluations were used to quantitate the levels of mRNA encoding PD-L1 and Chi3l1 and PD-L1 proteins in lungs from WT mice challenged with B16 cells (B16 +) or its PBS vehicle control (B16-) that were treated with Poly(I:C) or its vehicle control. (C) RT-PCR was used to quantitate the levels of mRNA encoding LAG3 in the lungs from WT mice that were treated intravenously with vehicle (B16 -) or B16 cells (B16 +) and randomized to receive poly(I:C) or vehicle control. (D) FACS evaluations of PD-L1 on CD68+ macrophages from lungs from WT mice that received B16 cells or their PBS controls and were treated with Poly(I:C) or its vehicle control. (E-F) RT-PCR was used to quantitate the levels of mRNA encoding PD-L1 (E) and LAG-3 (F) in the lungs from WT mice and Chi3l1 Tg+ mice that were treated intravenously with vehicle (B16 -) or B16 cells (B16 +) and randomized to receive poly(I:C) or vehicle control. Each dot represents an evaluation in an individual animal. The plotted values in panels A, C and D represent the mean±SEM of the evaluations represented by the individual dots. Panels B and D are representative of at least 2 similar evaluations. **P<0.01.

### Anti-Chi3l1 and anti-PD-1 interact to augment antitumor responses in melanoma lung metastasis

Since Chi3l1 stimulates PD-L1 and other ICPIs, studies were undertaken to determine if anti-Chi3l1 and anti-PD-1 interact in inducing antitumor responses. In these experiments, mice were treated with the antibodies individually and in combination. As can be seen in Fig. 7, FRG and anti-PD-1 individually inhibited melanoma metastasis in a dose dependent manner when compared to isotype controls (Fig. 7, A and B). Importantly, when mice were treated with 50 μg doses of the two antibodies the antitumor responses induced by the antibodies in combination exceeded the effects that were seen when the antibodies were used individually (Fig. 7, A and B). These effects appeared to be at least additive in nature. They demonstrate that anti-Chi3l1 and anti-PD-1 interact to augment antitumor responses in lung melanoma metastasis.

**Fig. 7.**
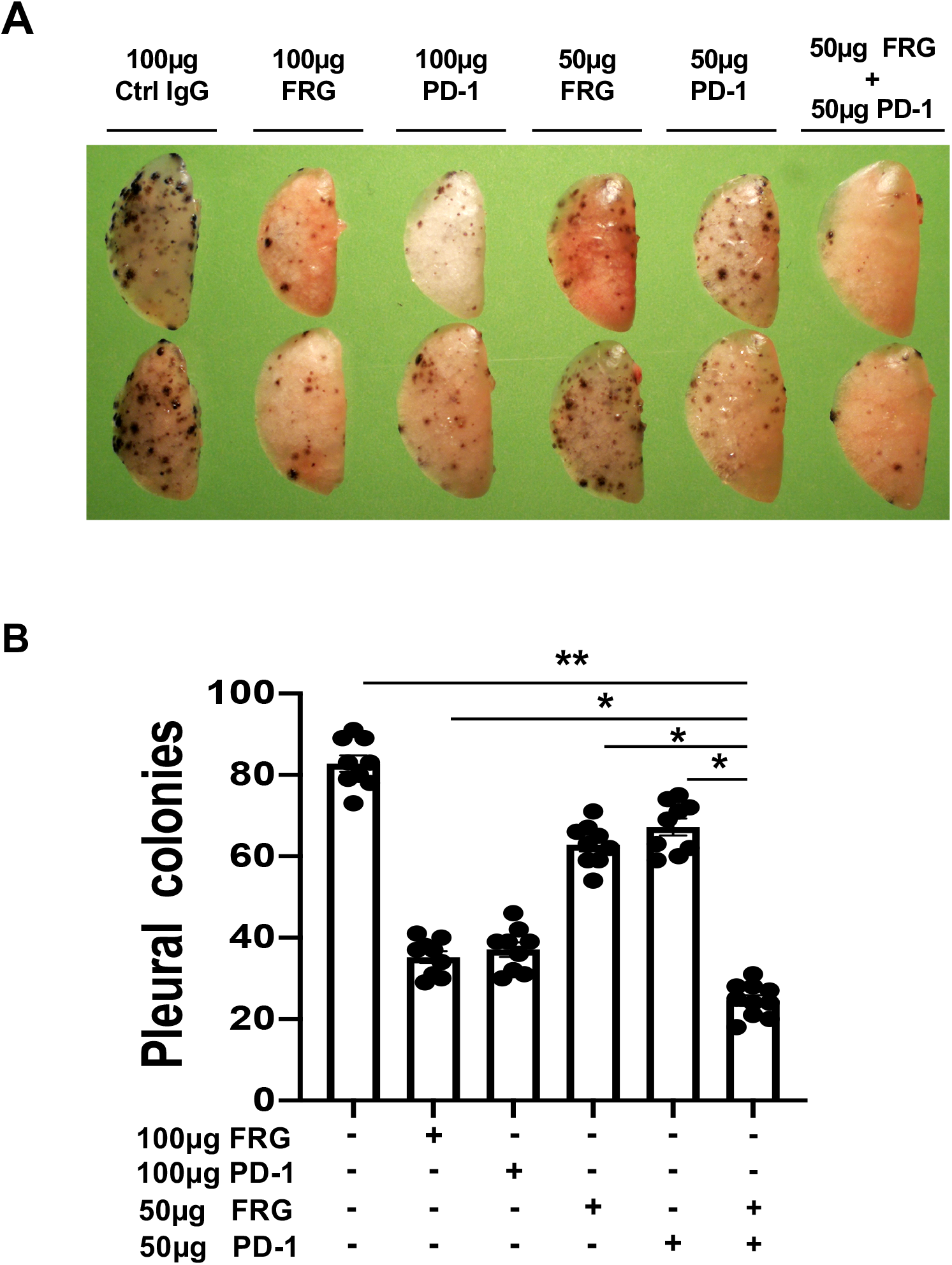
Anti-Chi3l1 and anti-PD-1 interact to augment antitumor responses in melanoma lung metastasis. WT mice were given B16-F10 (B16) melanoma cells or control vehicle and treated with control IgG, FRG and or anti-PD-1 antibodies, alone or in combination, and melanoma lung metastasis was evaluated 2 weeks later. (A) Representative lungs from mice treated with control IgG, FRG and or anti-PD-1 antibodies, alone or in combination. As noted, the antibodies were given at doses of 100μg or 50 μg every other day by intra-peritoneal injection. (B) The number of pleural melanoma colonies was quantitated in the lungs from the mice in panel A. Each dot is representative of an individual animal. Panel A is a representative of at least 3 similar evaluations. The values in panel B represent the mean ± SEM of the evaluations represented by the individual dots in the lungs from the experiment representatively illustrated in panel A. *P<0.05. **P<0.01.

### Antibodies that simultaneously target Chi3l1 and PD-1 induce CTL cell differentiation, PTEN expression and tumor cell death

In recent years it has become clear that combination therapy with ICP blockers can induce particularly potent responses in a variety of tumors including lung cancers (46–48). The studies noted above suggest that antibodies against Chi3l1 and PD-1 interact to enhance antitumor responses in melanoma metastasis. Thus, to further understand this interaction we compared the effects of FRG and anti-PD-1, alone and in combination, in a co-culture system containing activated T cells and A357 human melanoma cells. In these co-culture experiments, FRG and anti-PD-1 individually caused a significant increase in tumor cell apoptosis (Fig. 8A). When FRG and anti-PD-1 were administered simultaneously an additional increase in tumor cell death was seen. This effect appeared to be at least additive in nature (Fig. 8A). In all cases, the tumor cell death that was seen appeared to be mediated by cytotoxic T cells because FRG and anti-PD-1, alone and in combination, heightened T cell expression of perforin and granzyme (Fig. 8, B-C). Surprisingly, FRG and anti-PD-1, alone and in combination, also heightened the expression of the tumor suppressor PTEN (Fig. 8D). Identical results were seen in experiments that employed these antibodies and murine B16-F10 melanoma cell line (Fig. S5, A-D). When viewed in combination, these studies demonstrate that antibodies that simultaneously target Chi3l1 and PD-1 have impressive anti-tumor effects that are mediated by its ability to induce cytotoxic T cells and enhance tumor cell expression of PTEN.

**Fig. 8.**
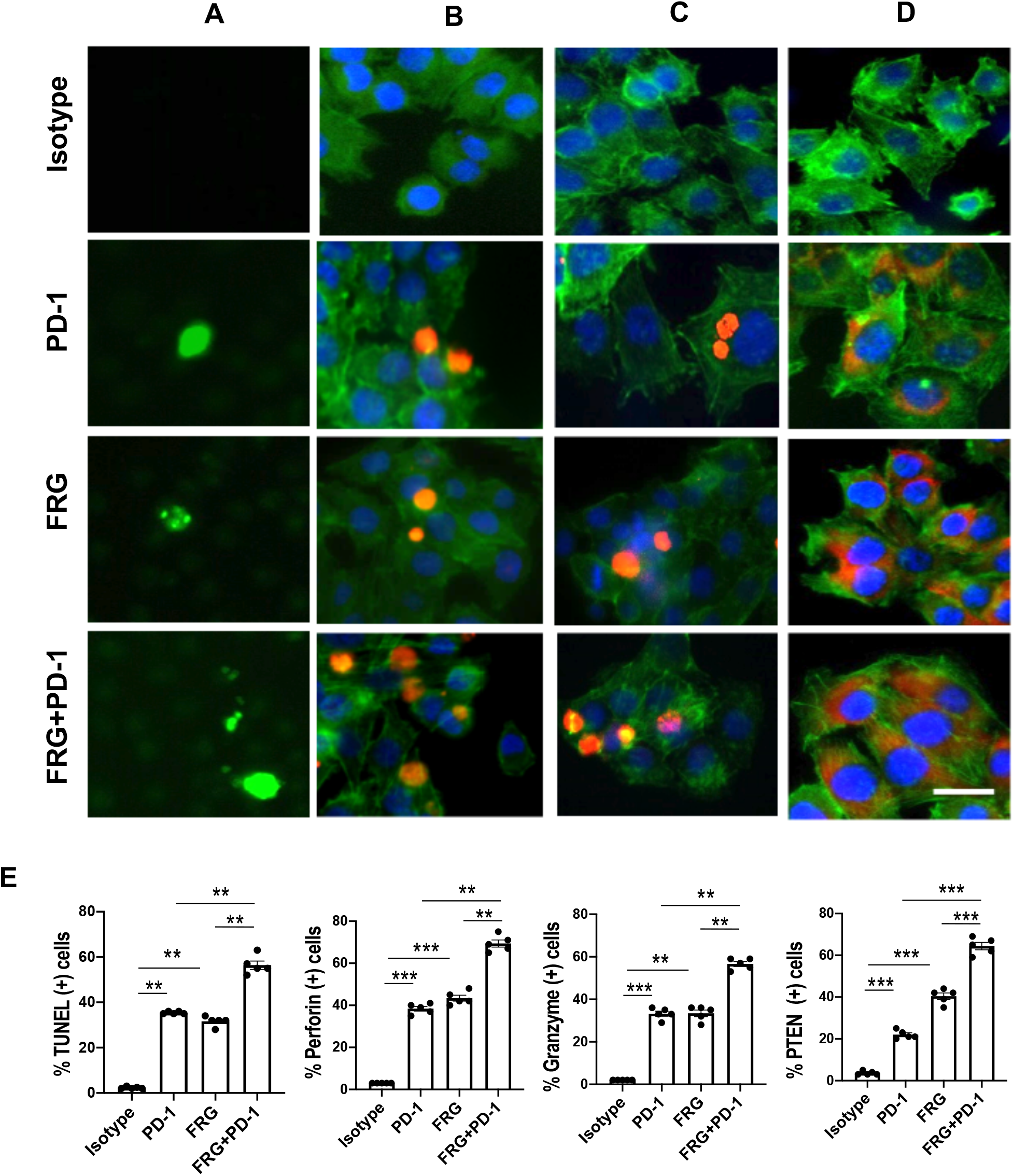
Antibodies that target Chi3l1 and PD-1 induce CTL-mediated tumor cell death responses and tumor cell PTEN expression. The antitumor effects of antibodies against Chi3l1 and PD-1 were evaluated in a co-culture system containing Jurkat cells and A375 human melanoma cells. Jurkat cells were activated by pretreatment with anti-CD3 and anti-CD28 (1 μg/ml each and incubated for 2 hrs in 5% CO2 and air at 37°C). The Jurkat cells were then cocultured with A357 human melanoma cells for 24 hours. These co-cultures were undertaken in the presence of the following antibodies; isotype control antibody (5μg/ml), anti-PD-1 or anti-Chi3l1 (FRG) alone (5μg/ml), or anti-Chi3l1 and anti-PD-1 in combination (2.5 μg/ml each) as noted. (Column A) Representative demonstration and quantitation of apoptotic tumor cell death using in situ cell detection kit -fluorescein dUTP. TUNEL (+) cells stain green. (Columns B-C) Representative demonstration and quantification of Jurkat T cell expression of perforin (B) and granzyme (C). Tumor cells are green and positive staining Jurkat cells are yellow-orange. (D) Representative demonstration and quantification of tumor cell PTEN. Tumor cells are green and PTEN is yellow-orange. (Row E) Quantification of the evaluations in Columns A-D. The % of TUNEL + tumor cells (Column A), % of Jurkat cells expressing perforin (column B) and granzyme (column C) and % of tumor cells expressing PTEN (column D) are illustrated. These evaluations were done using fluorescent microscopy (x20 of original magnification). In these quantifications, 10 randomly selected fields were evaluated. The values in panel F are the mean ± SEM of the noted 4 evaluations. **P<0.01. ***P<0.001. Scale bar=50μm, applies to all subpanels of A-E.

## DISCUSSION

The immune surveillance hypothesis proposed in the 1950’s that the immune system could recognize and reject cancer cells as foreign, in the same way that it reacts to microbes and transplanted organs (49). It has become clear over the last two decades that there is validity to this hypothesis and that tumors become shielded from immune elimination by aberrantly expressing ligands that normally interact with inhibitory immune receptors that protect “self” (49). To further understand the mechanisms that mediate these immunosuppressive responses, we tested the hypothesis that Chi3l1 plays a critical role in these tumor inhibitory events. These studies highlighted the ability of pulmonary metastasis to stimulate components of the PD-1-PD-L1 axis and other ICP molecules. Importantly, they also highlighted a novel relationship between Chi3l1 and ICPs by demonstrating that Chi3l1 stimulates PD-L1, PD-1, PD-L2 and other ICPs *in vivo* and *in vitro.* In the former, null mutations of or treatment with antibodies that target Chi3l1abrogated the induction of PD-L1 and other ICPs by metastatic tumors. Interestingly, transgenic Chi3l1, in the absence of tumor, reproduced these events, demonstrating that the effects of Chi3l1 are, at least partially, tumor-independent. In the latter, Chi3l1 was shown to stimulate macrophage PD-L1 and IFN-γ stimulation of PD-L1 was shown to be mediated, in part, by its ability to stimulate Chi3l1. These findings provide a potential mechanism(s) for the immunosuppressive effects of Chi3l1 that have been described in the context of breast cancer (50). They also describe a potentially important mechanism by which type I immune responses can be coopted to shut down antitumor immune responses (4, 5). These studies also have important therapeutic implications because they demonstrate that the simultaneous administration of anti-Chi3l1 and anti-PD-1 antibodies generate, at least additive, antitumor responses *in vivo* and that bispecific antibodies that simultaneously target Chi3l1 and PD-1 synergistically induce cytotoxic T lymphocyte differentiation and tumor cell death.

Early concepts of tumor immunity highlighted the importance of tumor cell expression of ligands like PD-L1 which bind to T cell PD-1 and suppress T cell activation (51). However, it is now clear that tumor infiltrating immune cells can be the most common cells expressing PD-L1 in tumor specimens (5) and that tumor cell expression of PD-L1 does not always predict patient responsiveness to PD-1/PD-L1-based therapeutic interventions (5). In fact, in some settings the levels of PD-L1 on immune cells are a better predictor of responsiveness to antiPD-1 than the levels of PD-L1 on tumor cells (5). In accord with this complexity, our studies demonstrate that metastatic B16-F10 cells and transgenic Chi3l1 stimulate PD-L1 in a variety of non-tumor pulmonary cells. Macrophages were particularly prominent in these responses and Chi3l1 was a powerful stimulator of macrophage PD-L1 in the murine lung and cultured bone marrow-derived macrophages. Studies of tumor-associated macrophages have demonstrated that some have a M1 (classic activation) phenotype. However, the majority appear to have a M2 (alternatively activated) phenotype with M2 cells predominating when the tumor metastasizes (52, 53). Previous studies from our laboratory and others have demonstrated that Chi3l1 contributes to macrophage diversity via the induction of M2 differentiation (12, 13, 50). PD-L1 is now known to also play a regulatory role in M1/M2 polarization with PD-L1 suppression decreasing M2 and augmenting M1 differentiation (52). When viewed in combination, these studies allow for the overall hypothesis that Chi3l1 stimulates macrophage PD-L1 which augments the accumulation of M2 macrophages and generates a microenvironment that fosters tumor growth and metastasis.

IFN-γ is a multifunctional type I cytokine that is known to induce antitumor responses via a number of mechanisms including the activation of macrophages and monocytes (44). In accord with this concept, the secretion of IFN-γ by stimulated blood mononuclear cells from patients with advanced cancer has been reported to be significantly decreased compared to cells from healthy controls (44, 54). In contrast, in certain settings, IFN-γ also acts to induce tumor progression (44). Although the mechanisms that underlie the tumor permissive effects of IFN-γ have not been fully defined, recent studies suggest that IFN-γ does this, in part, by inducing immune escape via the induction of PD-L1 (44). Our studies support this concept by demonstrating that IFN-γ stimulates PD-L1 in macrophages and other cells. They also demonstrate that IFN-γ stimulation of Chi3l1 is a critical event in this PD-L1 inductive response. To enhance the antitumor effects of IFN-γ it has been suggested that rIFN-γ be administered to patients with low IFN-γ activity and that anti-PD-1/L1 be co-administered in these circumstances (44, 54). Our studies add to this concept by suggesting that patients that are being treated with IFN-γ or that have high levels of IFN-γ activity could be treated with Chi3l1 inhibitors or FRG to maximize the therapeutic efficacy of IFN-γ. Further experimentation will be required to address these possibilities.

Because Chi3l1 lacks enzymatic activity when compared to true chitinases, studies were undertaken to determine if it mediated its functions via novel receptors. These studies demonstrated that Chi3l1 binds to, signals, and confers tissue responses via IL-13Rα2 and CRTH2 (15, 20). They also demonstrated that IL-13Rα2 plays a central role in a multimeric receptor complex called the chitosome that has an IL-13Rα2 alpha subunit and at least one beta subunit called TMEM219 (TMEM) (15, 18). The other beta subunit may be CD44 (variant 3) which is now known to bind IL-13Rα2 (55). These studies also demonstrated that Chi3l1 activates the MAPK, Akt/ Protein kinase B and Wnt/β-catenin signaling pathways with optimal MAPK and Akt activation requiring IL-13Rα2 and TMEM and Wnt/β-catenin signaling being mediated via a TMEM-independent mechanism (15, 18). To further understand the mechanisms by which Chi3l1 regulates PD-L1, we compared the ability of Chi3l1 to stimulate PD-L1 in macrophages from wild type mice, mice with null mutations of IL-13Rα2 or TMEM and wild type cells treated with a specific CRTH2 inhibitor. These studies demonstrate that IL-13Rα2 plays a critical role in Chi3l1 stimulation of macrophage PD-L1. In contrast, Chi3l1 induction of PD-L1 was not altered by null mutations of TMEM or by treatment with the CRTH2 inhibitor. These observations raise the interesting possibility that Wnt/β-catenin pathway activation plays a major role in this Chi3l1 stimulatory event. We cannot, however, rule out contributions from MAPK or Akt signaling because they have been implicated in the regulation of PD-L1 in other circumstances.

Activation of the RIG-I/RLH signaling pathway in tumors and the host microenvironment has recently been demonstrated to be a critical component of immune checkpoint blockade (ICB) induced by antibodies against CTLA4, alone and in combination with antibodies against PD-1 (42). When RIG-I/RLH innate immunity is appropriately induced ICB triggers caspase-3-mediated tumor cell death, cross presentation of tumor associated antigen by CD103+ dendritic cells and the accumulation of antigen specific CD8+ infiltrating T cells (42). It was speculated that nucleic acids leaking from disintegrating tumor cells during RIG-I-induced cell death are engulfed by host myeloid cells to induce RLH activation and type I IFN production (42). We previously demonstrated that RLH activation inhibits Chi3l1 production (41). The present studies demonstrate that RLH inhibition of Chi3l1, in turn, inhibits PD-L1 and other ICPs. They also demonstrate that, at least additive, antitumor effects are seen when mice with B16-F10 melanoma metastasis are treated simultaneously with anti-Chi3l1 and anti-PD-1 individually. These studies provide an additional mechanism that can explain how RIG-I/RLH activation fosters ICB antitumor responses. Specifically, they suggest that the antitumor effects of RIG-I/RLH activation are due to its ability to inhibit Chi3l1 and that Chi3l1 is a critical regulator of ICB. This raises the exciting possibility that interventions that inhibit or block Chi3l1 can be used to maximize the therapeutic efficacy of ICB.

LAG3 was cloned in 1990 and shown to have 20% homology with CD4 and bind to and inhibit T cell activation via MHC class II. It is upregulated as a feedback mechanism on activated and exhausted T cells (4). Our studies define, for the first time, a relationship between LAG 3 and Chi3l1 by demonstrating that Chi3l1 is a potent stimulator of LAG3. In so doing they demonstrate that the effects of Chi3l1 on ICPs are not specific for components of the PD-1-PD-L1 pathway. They also provide a potential explanation for the synergy between interventions that target Chi3l1 and PD-1 because LAG3 and PD-1 are known to interact synergistically to suppress T cell activation (4, 56).

Lung cancer is the leading cause of cancer-related mortality worldwide (57). In keeping with its importance, our knowledge of tumor immunology and ICP inhibitors has led to the development of a number of new therapeutics that have been assessed in metastatic and primary lung cancers (49). In contrast to the prior therapeutic dogma that focused initially on tumor resection followed by chemotherapy and or radiation therapy, ICP inhibitors are now first or second line therapeutics (58). When these agents are employed remarkable responses are seen in some individuals. However, overall, only a minority of patients respond to these therapies and the responses that are seen are often not durable (59). To address these issues combination immunotherapy has been attempted. The simultaneous treatment with anti-PD-1 and anti-CTLA4 has been most commonly employed (60). This combination has engendered interesting effects is in some malignancies. However, it has not proven successful in lung cancers. In addition, its use is limited by toxicities such as the pneumonitis that is caused by the exaggerated immune response that the antibody combination induces (60). In keeping with the concept that combination immunotherapy may be therapeutically useful, we evaluated the effects of anti-Chi3l1 (FRG) and anti-PD-1, alone and in combination. These studies demonstrate that the simultaneous treatment with antibodies that target Chi3l1 and PD-1 individually results in additive antitumor responses. Based on our finding that RLH activation inhibits Chi3l1 we can also envision similarly augmented anticancer response if RLH activators and antiPD-1 are coadministered. Whether the safety profile of these immunotherapeutic combinations is better or worse than anti-PD-1 plus anti-CTLA4 will need to determined.

To further understand the mechanisms that underlie the enhanced antitumor effects of anti-Chi3l1 and anti-PD-1 their effects in a T cell-melanoma cell coculture system were evaluated. These studies demonstrate that FRG and anti-PD-1 are effective inducers of tumor cell death. They also demonstrated that these antitumor responses were augmented when the antibodies were used in combination. Lastly, they demonstrated that the tumor cell death responses were due to the ability of FRG and anti-PD-1 to induce perforin +, and granzyme+ cytotoxic T cells. These findings agree with a fundamental principle of tumor immunology, that cancer cells can be eliminated by host cytotoxic T cells (61). The heightened nature of the responses induced by FRG an anti-PD-1 in combination further support our contention that multiple different immunoregulatory pathways are involved in this response. In addition to the ability of anti-PD-1 to block the interactions of PD-1 with PD-L1 and/or PD-L2, interventions that diminish the production and or effector functions of Chi3l1 inhibit the ability of Chi3l1 to inhibit cell death (apoptosis and pyroptosis) and augment the accumulation of IFNα/β, chemerin and its receptor ChemR23, phosphorylated coffilin and LimK2 (41). Chi3l1-based interventions also augment the expression of the tumor suppressor PTEN, and decrease M2 macrophage differentiation and augment type I immune responses (41). When viewed in combination, these studies demonstrate that antibodies that simultaneously targeting Chi3l1 and PD-1 cause augmented tumor suppressor gene activation, CTL differentiation, and tumor cell death. When viewed in combination, these findings highlight the potentially exciting therapeutic consequences that come from a heightened understanding of the relationships between Chi3l1 and ICPI.

In conclusion, these studies demonstrate that Chi3l1 stimulates PD-L1 and other ICP moieties such as LAG3 *in vivo* and *in vitro.* They also demonstrate that the simultaneous treatment of pulmonary metastatic melanoma with individual antibodies against Chi3l1 and PD-1 generates an augmented antitumor response. These findings predict that interventions that simultaneously target Chi3l1 can augment the efficacy of immune checkpoint blockade in response to anti-PD-1 in lung cancer and potentially other malignancies. Additional investigations of the biology and therapeutic consequences of interactions between Chi3l1 and immune checkpoint inhibitors are warranted.

## MATERIALS AND METHODS

### Genetically modified mice

Mice with null mutations of Chi3l1 (*Chi3l1^-/-^*) and TMEM219 (*TMEM219^-/-^*), and transgenic mice in which Chi3l1 was targeted to the lung with the CC10 promoter (*Chi3l1 Tg*) have been generated and characterized by our laboratory as previously described (13, 18). Mice with null mutations of IL-13Rα2 (*IL-13Rα2^-/-^*) mice were a generous gift from Dr. Grusby (62) and have been backcrossed for more than 10 generations onto a C57/BL6 background. Gal3 null mutant mice (*Gal3^-/-^*) on a C57/BL6 background and Wild-Type C57BL/6J mice were purchased from the Jackson Laboratories (Bar Harbor, Me USA). These mice were between 6-12 weeks old when used in these studies. All animals were humanely anesthetized with Ketamine/Xylazine (100mg/10mg/kg/mouse) before any intervention. The protocols that were used in these studies were evaluated and approved by the Institutional Animal Care and Use Committee (IACUC) at Brown University.

### Western blot analysis

Protein lysates from macrophages and whole mouse lungs were prepared with RIPA lysis buffer (ThermoFisher Scientific, Waltham, MA, USA) containing protease inhibitor cocktail (ThermoFisher Scientific) as per the manufacturer’s instructions. 20 to 30 μg of lysate protein was subjected to electrophoresis on a 4–15% gradient mini-Protean TGX gel (Bio-rad, Hercules, CA, USA). It was then transferred to a membrane using a semidry method with a Trans-Blot Turbo Transfer System (Bio-rad). Membranes were blocked with Tris-buffered saline with Tween20 (TBST) with 5% non-fat milk for 1 hour at room temperature. After blocking, the membranes were incubated with the primary antibodies overnight at 4^0^ in TBST and 5% BSA. The membranes were then washed 3 times with TBST and incubated with secondary antibodies in TBST, 5% non-fat milk for 1 hour at room temperature. After 3 additional TBST washes, Supersignal West Femto Maximum Sensitivity Substrate Solution (Thermofisher Scientific) was added to the membrane and immunoreactive bands were detected by using a ChemiDoc (Bio-Rad) imaging system.

### RNA extraction and Real-time qPCR

Total cellular RNA was obtained using TRIzol reagent (ThermoFisher Scientific) followed by RNA extraction using RNeasy Mini Kit (Qiagen, Germantown, MD) according to the manufacturer’s instructions. mRNA was measured and used for real time (RT)-PCR as described previously (13, 17). The primer sequences used in these studies are summarized in Table S1. Ct values of the test genes were normalized to the internal housekeeping gene β-actin.

### Generation of α-Chi3l1 and chimeric bispecific antibodies against Chi3l1 and PD-1

The murine monoclonal anti-Chi3l1 antibody (FRG) was generated using peptide antigen (FRGQEDASPDRF, amino acid 223-234 of human Chi3l1) as immunogen. This monoclonal antibody specifically detects both human and mouse Chi3l1 with high affinity (kd ≈1.1×10^-9^). HEK-293T cells were transfected with the anti-Chi3l1 construct using Lipofectamine™ 3000 (Invitrogen, L3000015, CA). Supernatant was collected for 7 days and the antibody was purified using a protein A column (ThermoFisher Scientific, 89960; IL). The original murine anti-PD1 monoclonal antibody (63) was a generous gift from Prof. Lieping Chen (Medical Oncology, Yale School of Medicine).

### Melanoma lung metastasis and antibody treatment

B16-F10, a mouse melanoma cell line, was purchased from ATCC (Cat#: CRL-6475, Manassas, VA) and maintained in Dulbecco’s Modified Eagles Medium (DMEM) supplemented with 10% Fetal bovine Serum (FBS) and 1% penicillin & streptomycin. When the cells formed an 80% confluent monolayer, they were collected, adjusted to the concentration of 10^6^ cells/ml and injected into the mice via their lateral tail veins (2×10^5^ cells/mouse in 200 μl of DMEM). As previously described, intraperitoneal injection of the noted doses of anti-Chi3l1 (FRG) and anti-PD-1, alone and in combination and isotype control IgG (IgG2b) were started on the day of the B16 tumor cell challenge and continued every other day for 2 weeks (40) (41). Metastasis was assessed and quantified by counting the melanoma colonies (black dots) on the pleural surface as previously described (40, 41).

### Fluorescence-activated cell sorting (FACS) analysis

Single cell suspensions from whole mouse lungs were prepared using the lung dissociation kit (Miltenyi Biotec, Auburn, CA) as per the manufacturer’s instructions. Cells were stained with fluorescently labeled antibodies directed against PD-L1, PD-L2 (*InVivoMAb;* BioXcell, West Lebanon, NH), CC10, SPC (Santa Cruz Biotechnology, Santa Cruz, CA), CD3, CD68-PE, CD11b-FITC (BD Pharmingen, Billerica, MA, USA), PD-L1-PE-cy7, and F4/80-FITC (Biolegend, Dedham, MA). Flow cytometry data was collected using the BD FACSAriaIIIu and analyzed with FlowJo V10 software.

### Immunohistochemistry

Formalin-fixed paraffin embedded (FFPE) lung tissue blocks were serially sectioned at 5 μm-thickness and mounted on glass slides. After deparaffinization and dehydration, heat-induced epitope retrieval was performed by boiling the samples in a steamer for 30 minutes in antigen unmasking solution (Abcam, antigen retrieval buffer, 100x citrate buffer pH:6.0). To prevent nonspecific protein binding, all sections were blocked in a ready-to-use serum free protein blocking solution (Dako/Agilent, Santa Clara, CA) for 10 minutes at room temperature. The sections were then incubated with primary antibodies (a-PD-L1, a-CC10, a-SPC, a-CD68) overnight at 4°C. After three washings, fluorescence-labeled secondary antibodies were incubated for 1 hour at room temperature. The sections were then counterstained with DAPI and cover slips were added.

### Double label immunohistochemistry

Double label immunohistochemistry was employed as previously described by our lab (18).

### Macrophage preparation and treatment

Murine peritoneal and bone marrow derived macrophages (BMDM) were prepared as previously described (64). They were then incubated with the noted concentrations of recombinant mouse (rm) Chi3l1 or IFNγ (R&D Systems, Minneapolis, MN) or CAY10471 (Cayman Chemical, Ann Arbor, MI). After incubating for the desired periods of time the macrophages were collected and evaluated using RT-PCR, Western blotting or flow cytometric assays as noted above.

### T cell culture and activation

Jurkat T cells (1×10^5^) were grown in complete RPMI (10% FBS and 1%Penicillin and streptomycin). They were then activated by incubating them with anti-human CD3 antibody (5μg/ml) (Biolegend) and anti-human CD28 antibody (5μg/ml) (Millipore #MABF408, Burlingtone, MA) simultaneously for 2 hours in 5% CO2 and air at 37°C. After the incubation, the cells were washed twice to remove the extra antibody.

### Melanoma cells and Jurkat cell co-cultures

Human melanoma A375 cell line (# CRL-1619) and B16-F10 mouse melanoma cells (#CRL-6475) were purchased from ATCC (Manassas, VA) and cultured according to the instructions provided by the vendor. The Jurkat cells (1×10^5^) were cultured and activated as described above. After incubation, the cells were washed and B16 melanoma and activated Jurkat cells were resuspended together at a 1: 6 ratio in complete RPMI media, dispensed into the multi-well slide chambers and incubated in 5% CO2 and air at 37°C. One hour later, the co-cultured cells were treated with the isotype control antibody or test antibodies. The effects of the isotype control antibodies were compared to the effects of antibodies against Chi3l1 (FRG; 5μg), PD-1 (5μg), and FRG+PD-1 administered simultaneously (2.5μg each). After incubation for an additional 48 hours, cells were subjected to TUNEL evaluations and immunofluorescence evaluations of perforin, granzyme-B and PTEN by as described. below.

### Measurement of cellular apoptosis and cytotoxic cell death responses

TUNEL staining using fluorescein-labeled dUTP was employed to assess apoptosis and cytotoxic cell death responses. After 48 hour incubation with test and control antibodies, cocultured cells were fixed in the 4% paraformaldehyde in PBS, permeabilized and blocked. In between each step, the cells were washed twice with PBS (1X). The cells were then stained with the in-situ cell death detection kit, fluorescein (Roche, Mannheim, Germany) as per the manufacturer’s instructions.

### Immunofluorescence staining

To evaluate the activation and differentiation of cocultured T cells and expression of the tumor suppressor PTEN, immunofluorescence (IF) staining was carried out using antibodies against perforin (1:1000), granzyme-B (1:1000) (Biolegend) and PTEN (1:1000) (Cell signaling, Danvers, MA). After 48 hour incubation with test and control antibodies, cocultured cells were fixed in the 4% paraformaldehyde in PBS, permeabilized and blocked. Then the cells were incubated overnight at 4°C with primary antibodies noted above, washed twice with PBS (1X) and incubated for 2hours at 37°C with secondary detection antibodies (1:500) and phalloidin (1: 3000) (Invitrogen, Waltham, MA) for cytoskeleton staining. The cells were then washed, mounted with VECTASHIELD® antifade mounting medium (Vector Laboratories Inc.) and evaluated at 20X via fluorescence microscopy.

### Quantification and Statistical analysis

Statistical evaluations were undertaken with SPSS or GraphPad Prism software. As appropriate, groups were compared with 2-tailed Student’s *t* test or with nonparametric Mann-Whitney *U* test. Values are expressed as mean ± SEM. Statistical significance was defined as a level of *P* < 0.05.

## ACKNOWLEDGEMENT

This work was supported by National Institute of Health (NIH) grants U01 HL108638 (J.A.E.), PO1 HL114501(J.A.E.), and R01 HL115813 (C.G.L) from NHLBI and USAMRMC W81XWH-17-1-0196 (J.A.E) from Department of Defense.

## Author Contributions

Conception and design: BM, JAE, CGL, SK, BA; Data collection: BM, BA, SK, CML; Analysis and interpretation: BM, BA, SK, CML, JSK, CGL, JAE; Drafting the manuscript for important intellectual content: JAE, CGL

