## Supplemental Tables and Figures for "Chitinase 3-like-1 Stimulates PD-L1 and Other Immune Checkpoint Inhibitors"

### Supplementary Table

Table S1. Sequences of RT-PCR primers used in this study

| Gene | Sequence (5'to 3') | Length |
|---|---|---|
| PD-1-S | ACCCTGGTCATTCACTTGGG | 20 |
| PD-1-AS | CATTTGCTCCCTCTGACACTG | 21 |
| PD-L1-S | GCTCCAAAGGACTTGTACGTG | 21 |
| PD-L1-AS | TGATCTGAAGGGCAGCATTTC | 21 |
| PD-L2-S | CTGCCGATACTGAACCTGAGC | 21 |
| PD-L2-AS | GCGGTCAAAATCGCACTCC | 19 |
| LAG3-S | CTGGGACTGCTTTGGGAAG | 19 |
| LAG3-AS | GGTTGATGTTGCCAGATAACCC | 22 |
| Actin-S | GGCTGTATTCCCCTCCATCG | 20 |
| Actin-AS | CCAGTTGGTAACAATGCCATGT | 22 |

### Supplementary Figures

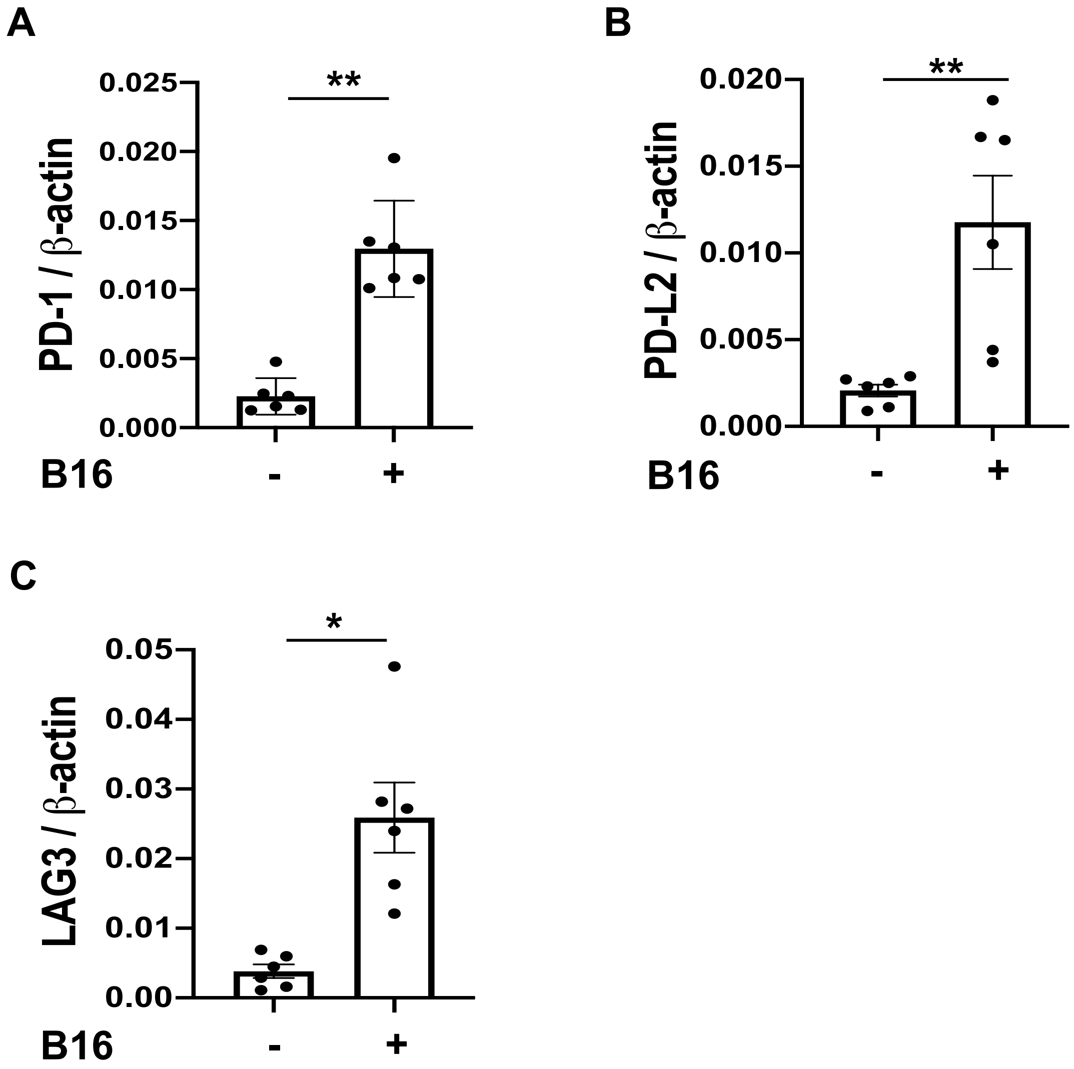


**Fig.S1. Pulmonary melanoma metastasis stimulates PD-1, PD-L2 and LAG3.** 8 week old WT mice were given B16-F10 (B16) melanoma cells or control vehicle (PBS) and the expression of immune checkpoint molecules was evaluated 2 weeks later. Real-time RT-PCR (RT-PCR) was used to quantitate the levels of mRNA encoding PD-1 (A), PD-L2 (B) and LAG3 (C) in the lungs from mice treated intravenously with PBS (B16 -) or B16 cells (B16 +). Each dot represents an evaluation in an individual animal. The plotted values represent the mean±SEM of the evaluations represented by the individual dots. *$p<0.05$, **$p<0.01$.

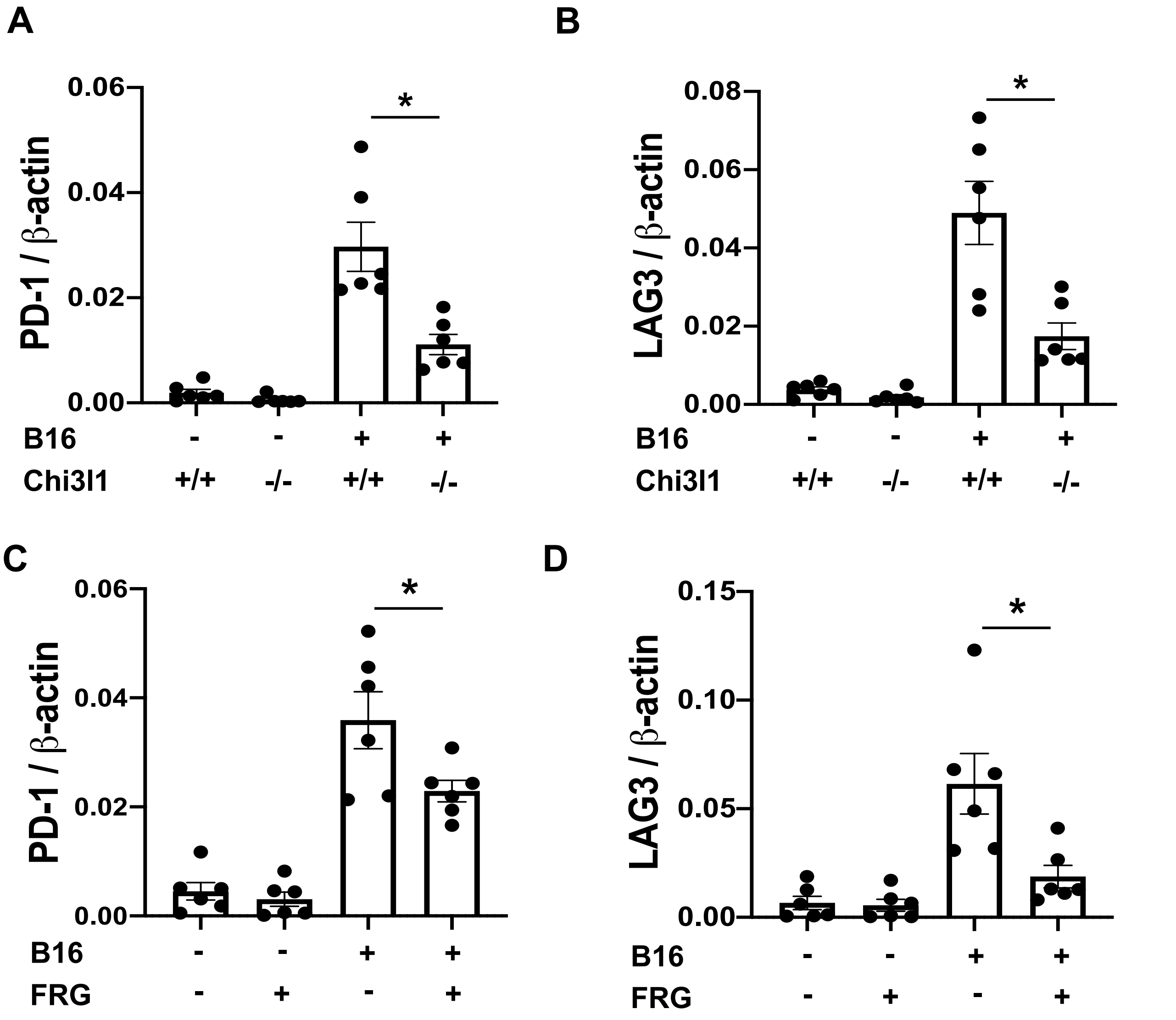


**Fig.S2. Chi3l1 plays a critical role in B16 melanoma stimulation of pulmonary PD-1 and LAG3.** 8 week old WT (+/+) and Chi3l1 null (-/-) (Chi3l1$^{-/-}$) mice were given B16 melanoma cells or vehicle control. They were then treated with an anti-Chi3l1 antibody (FRG) or isotype control antibodies and the expression of PD-1 or LAG3 was evaluated 2 weeks later. (A-B) RT-PCR was used to quantitate the levels of mRNA encoding PD-1 and LAG3 in the lungs from wild type (WT) (Chi3l1 +/+) and Chi3l1 null (Chi3l1 -/-) mice given PBS vehicle (B16 -) or B16 cells (B16 +). Each dot represents the evaluation in an individual animal. (C-D) RT-PCR evaluation of the levels of mRNA encoding PD-1 and LAG-3 in the lungs from WT mice treated with PBS vehicle (B16 -) or B16 cells (B16 +) that were treated with FRG (FRG+) or without FRG (FRG-). Each dot represents the evaluation in an individual animal. The plotted values represent the mean ± SEM of the noted evaluations represented by the individual dots. *$p<0.05$,

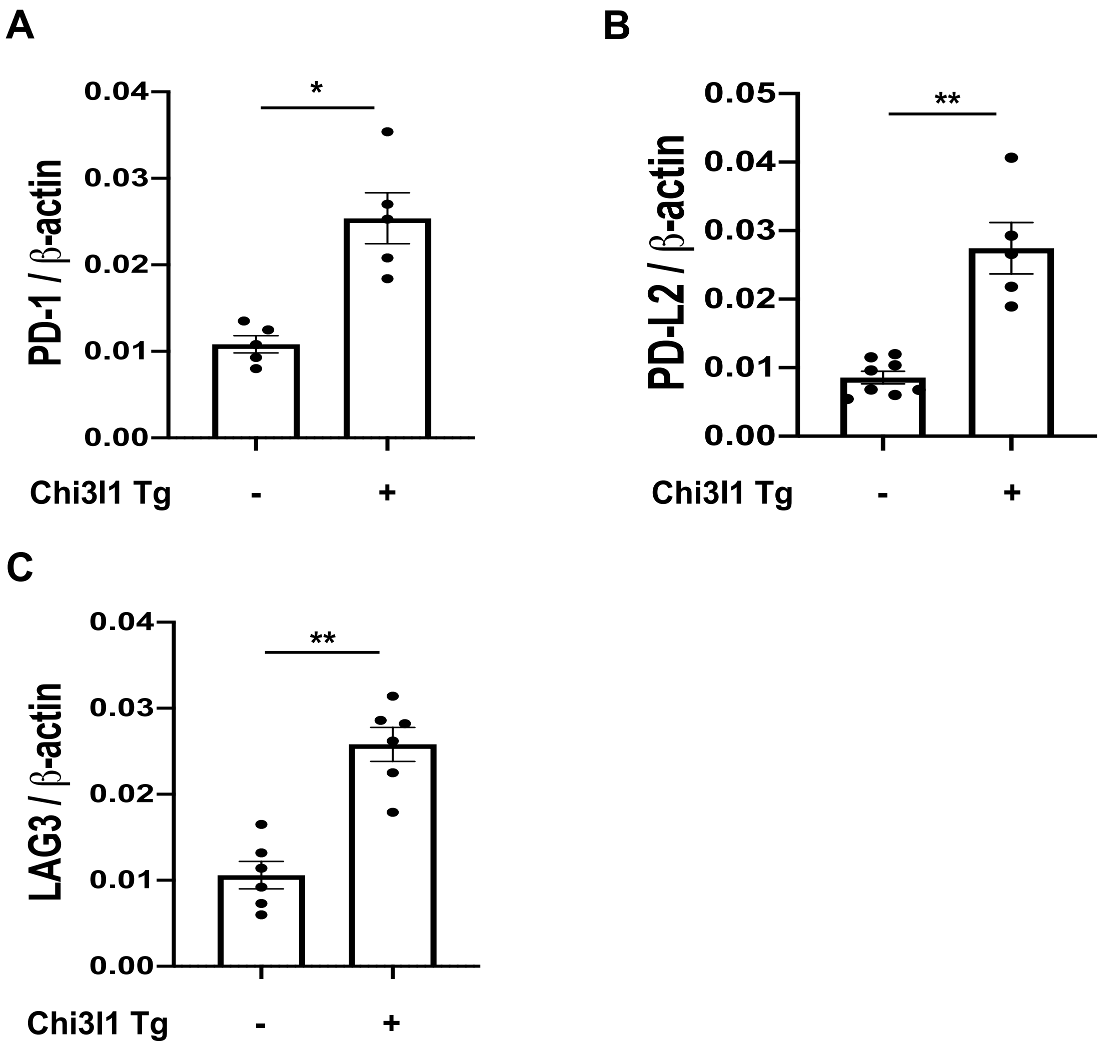


**Fig.S3. Transgenic Chi3l1 stimulates PD-1, PD-L2 and LAG3 in the normal lung.** 8 week old WT (-), Chi3l1 Tg (+) mice were used to evaluate the levels of mRNA encoding PD-1, PD-L2, and LAG3 in the lung. RT-PCR was used to quantitate the levels of mRNA encoding PD-1, PD-L2 and LAG3 in the lungs from WT mice (Chi3l1 Tg -) and mice in which Chi3l1 was overexpressed in the lung in a transgenic manner (Chi3l1 Tg +). Each dot represents the evaluation in an individual animal. The plotted values represent the mean ± SEM of the noted evaluations represented by the individual dots. *p<0.05,**p<0.01.

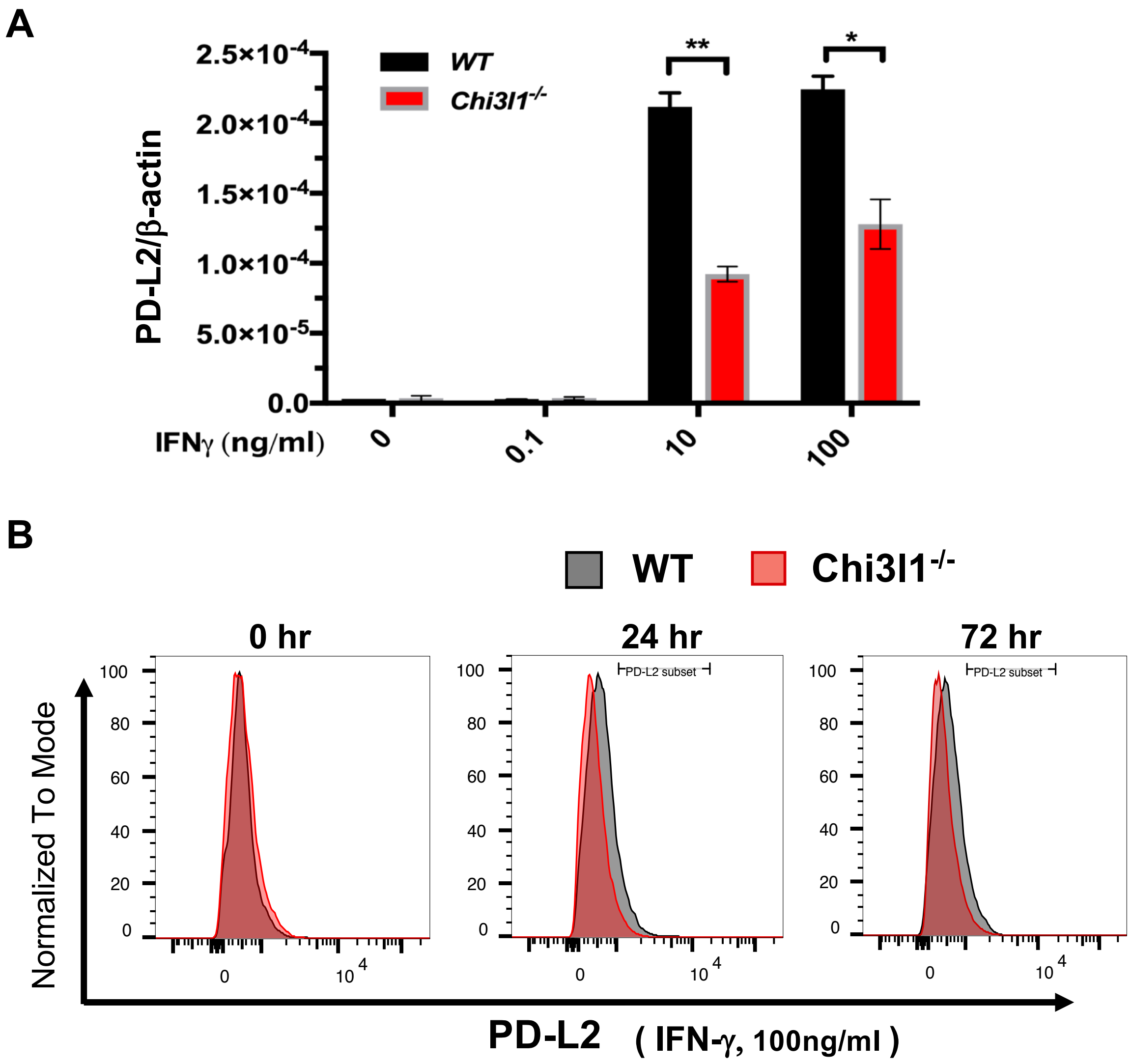


**Fig.S4. IFN-γ-stimulates macrophage PD-L2 via a Chi3l1-dependent mechanism.** Bone marrow derived macrophages (BMDM) prepared from 6-8 weeks old male WT and Chi3l1$^{-/-}$ mice and were used to evaluate the importance of Chi3l1 in rIFN-γ stimulation of PD-L2. (A) rIFN-γ-stimulation of PD-L2 mRNA expression in BMDM from WT and Chi3l1$^{-/-}$ mice. (B) FACS evaluations of the ability of rIFN-γ to stimulate PD-L1 in BMDM prepared from WT and Chi3l1$^{-/-}$ mice. The values in panel A represent the mean±SEM of the noted evaluations (n=5 mice/each). $^*p<0.05$, $^{**}p<0.01$. Panel B is a representative of a minimum of 2 separate evaluations.

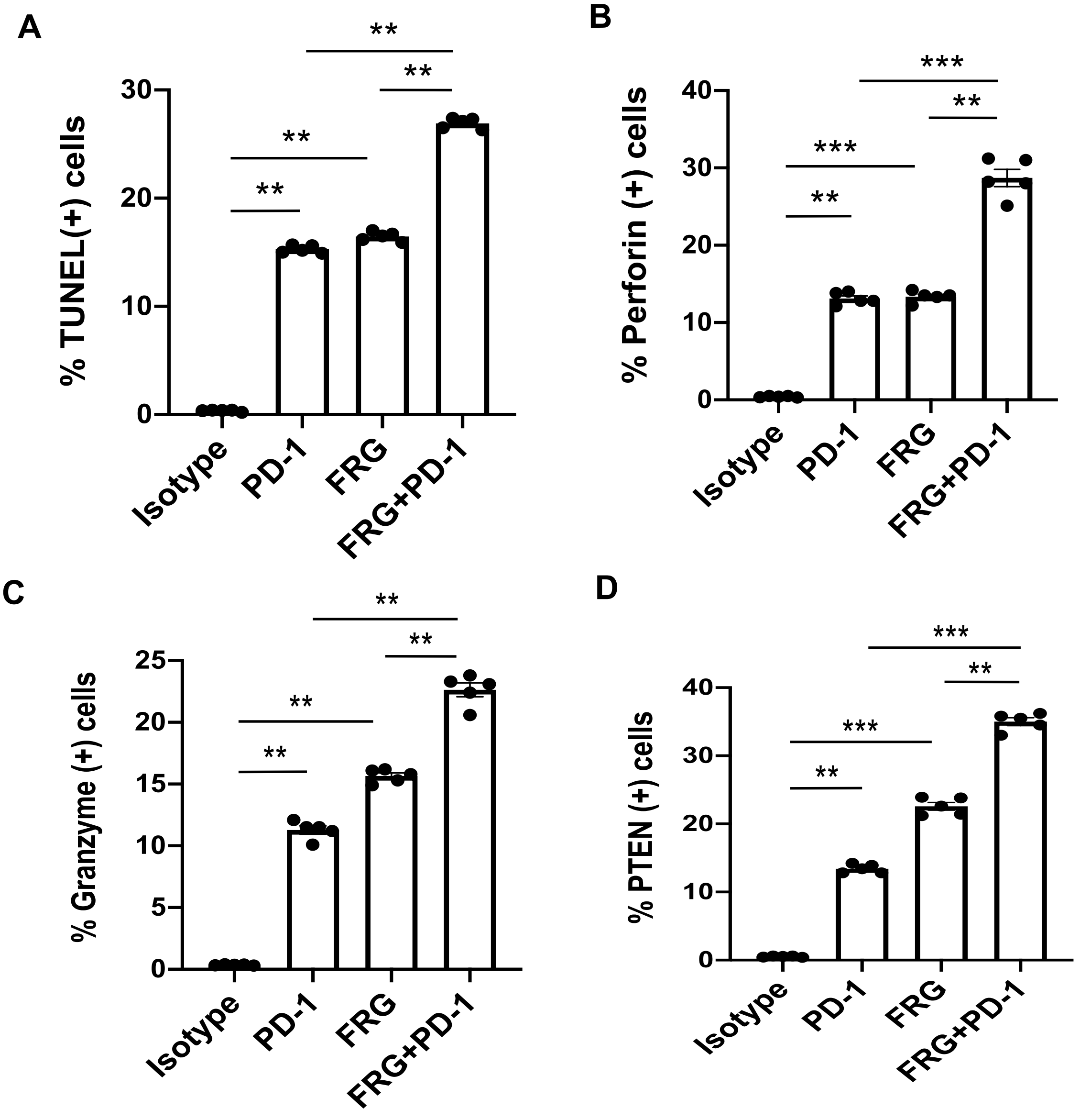


**Fig. S5: Effects of antiChi3l1 and anti-PD-1 on CTL-mediated tumor cell responses and PTEN accumulation**. The antitumor effects of anti-Chi3l1 and anti-PD-1 antibodies, alone and in combination, were tested in co-cultures of of Jurkat cells and B16-F10 murine melanoma cells as described in the Materials and Methods. (A) Quantitation of apoptotic tumor cell death using in situ cell death detection kit-fluorescein dUTP. Quantification of T cell perforin (B) and granzyme (C) accumulation. (D) quantification of tumor cell PTEN accumulation in co-cultures. These evaluations were undertaken using fluorescent microscopy (x20 of original magnification). In these quantifications, 10 randomly selected fields were evaluated. The values in in these panels are the mean ± SEM. **p<0.01, ***<0.001.
